# Data-Driven Image Analysis to Determine Antibody-Induced Dissociation of Cell-Cell Adhesion and Antibody Pathogenicity in Pemphigus Vulgaris

**DOI:** 10.1101/2024.10.09.617446

**Authors:** Amir Ostadi Moghaddam, Xiaowei Jin, Haiwei Zhai, Bahareh Tajvidi Safa, Kristina Seiffert-Sinha, Merced Leiker, Jordan Rosenbohm, Fanben Meng, Animesh A. Sinha, Ruiguo Yang

## Abstract

Pemphigus vulgaris (PV) is a blistering autoimmune disease that affects the skin and mucous membranes. The precise mechanisms by which PV antibodies induce a complete loss of cohesion of keratinocytes are not fully understood. But it is accepted that the process starts with antibody binding to desmosomal targets which leads to its disassembly and subsequent structural changes to cell-cell adhesions. In vitro immunofluorescence imaging of desmosome molecules has been used to characterize this initial phase, often qualitatively. However, there remains an untapped potential of image analysis in providing us more in-depth knowledge regarding ultrastructural changes after antibody binding. Currently, there is no such effort to establish a quantitative framework from immunofluorescence images in PV pathology. We take on this effort here in a comprehensive study to examine the effects of antibodies on key adhesion molecules and the cytoskeletal network, aiming to establish a correlation of ultrastructural changes in cell-cell adhesion with antibody pathogenicity. Specifically, we introduced a data-driven approach to quantitatively evaluate perturbations in adhesion molecules, including desmoglein 3, E-cadherin, as well as the cytoskeleton, following antibody treatment. We identify distinct immunofluorescence imaging signatures that mark the impact of antibody binding on the remodeling of the adhesion molecules and introduce a pathogenicity score to compare the relative effects of different antibodies. From this analysis, we showed that the biophysical response of keratinocytes to distinct PV associated antibodies is highly specific, allowing for accurate prediction of their pathogenicity. For instance, the high pathogenicity scores of the PVIgG and AK23 antibodies show strong agreement with their reported PV pathology. Our data-driven approach offers a more detailed framework for the action of autoantibodies in pemphigus and has the potential to pave the way for the development of effective novel diagnostic methods and therapeutic strategies.

**SIGNIFICANCE:** Pemphigus vulgaris (PV) presents a critical unmet medical challenge due to its autoimmune-induced disruption of skin cell adhesion. Our study presents a data-driven approach to quantitatively analyze changes in adhesion molecules and the cytoskeleton upon exposure to various PV antibodies. By introducing a pathogenicity score, we pinpoint the specific impacts of different antibodies on various proteins, build association among these antibodies, and reveal the contribution of previously overlooked non-desmosomal antibodies, broadening the understanding of PV pathology. Although centered on PV, our method offers a versatile framework applicable for evaluating the effects of other antibodies and drugs, paving the way for new diagnostic tools for personalized medicine.

## INTRODUCTION

Pemphigus vulgaris (PV) is a rare yet severe autoimmune disorder driven by the immune system’s aberrant production of autoantibodies against desmosomal proteins, primarily desmoglein (Dsg)-3 and 1 (1). Binding of these antibodies ultimately disrupts desmosomal adhesion in keratinocytes, induces acantholysis, and leads to blister formation by mechanisms that remain to be elucidated (2). The pathogenic potential of auto-antibodies has been estimated in vitro using keratinocyte dissociation assays (3,4) and by imaging disorganized adhesion molecules along the cell-cell contact by immunofluorescence (5,6). In general, keratinocytes exposed to PV antibodies exhibit fragmented Dsg3 staining, an increased distance between Dsg3 fluorescence peaks across neighboring cells, and decreased Dsg3 intensity at cell-cell contacts (7) due to Dsg3 internalization upon the initiation of desmosome disassembly. E-cadherin is also impacted in the PV condition either passively by the biophysical rearrangement of the cell-cell adhesion (8) or actively through signaling (9–11). Some reports even suggest that E-cadherin is a direct target of PV antibodies (12). In imaging, E-cadherin displays a distorted pattern to their tightly packed distribution along cell-cell contacts when keratinocytes are treated with PV antibodies (13).

Additionally, cytoskeleton reorganization represents a direct biophysical consequence once adhesion integrity is compromised. Indeed, keratin retraction plays a crucial role in PV pathogenesis. Desmosomes tether keratins to sites of intercellular adhesion, creating a cellular scaffold that imparts mechanical strength to tissues (14). Desmosome disassembly after antibody binding physically uncouples keratin from the desmosomal complex in response to cellular tension (15,16). Imaging techniques reveal that keratin retracts from the cell periphery as evident by the decreased number of keratin filaments running perpendicular to the cell membrane, and keratin filaments condensed to thicker bundles with an evident curvature change (17,18). Actin remodeling is another biophysical transformation induced by PV antibodies (19). It has been reported that PV antibody interferes with actin dynamics and destabilizes the junction associated actin belt (19,20). A loss of the peripheral actin band is often observed along with an enhanced cytoplasmic distribution of actin filaments (19). Actin dynamics are also heavily regulated by RhoA and P38 MARPK, molecules that are downstream of antibody binding (21).

Different PV-associated antibodies lead to varied changes in adhesion molecules and in cytoskeleton remodeling, indicative of their differential action mechanism and level of pathogenicity. For instance, PVIgG from patients can affect both Dsg3 and Dsg1 while the monoclonal antibody AK23 targets only Dsg3, leading to differences in biophysical remodeling at cell-cell adhesions. Recent studies have now also identified non-Dsg antibodies that are increased in patients vs. healthy controls and potentially contribute to PV pathology (22).

Image analysis of texture information, such as entropy, correlation, homogeneity, and contrast, has been used for the detection of subtle changes in tissues and cellular structures from biomedical images as a marker for disease progression. These practices have been performed on images collected from immunohistochemistry staining (23,24), MRI (25,26) and CT scanning (27). For instance, texture features were used to classify colon cancer cells from biological images. By employing parameters such as correlation, entropy, and contrast, normal and abnormal cells were distinguished with promising classification accuracy (28). These studies, along with the work of others (29–33), demonstrate the versatility and effectiveness of texture features in medical imaging applications for identifying and characterizing various biological conditions. In many cases, texture features are used as inputs for machine learning models, significantly enhancing the effectiveness in accurately classifying diseases (28,31).

Thus far there has been no such effort in the understanding of PV pathology using advanced image analysis. The images we collected documenting biophysical changes in combination with advanced image analysis offer an unprecedented level of detail regarding autoantibody induced biophysical transformations relevant to disease pathomechanisms. To establish a correlation of ultrastructural changes in cell-cell adhesion with antibody pathogenicity, we employed a data-driven approach to quantitatively evaluate immunofluorescence images collected from keratinocytes upon exposure to various PV antibodies. From these analyses, we identify distinct imaging features that mark the impact of antibody binding on the remodeling of the adhesion molecules and the cytoskeleton. These features show that the biophysical response of keratinocytes to different PV antibodies is highly specific, allowing for accurate prediction of antibody pathogenicity. Indeed, we introduced a *pathogenicity score* to compare the relative effects between different antibodies. The high pathogenicity scores of PVIgG and AK23 revealed in our studies show a strong agreement with their reported PV pathology in the literature. In addition to showing the relative strength of each antibody in terms of pathogenic disruption of cell-cell adhesion, our analysis also reveals correlative similarities among different antibodies. For instance, we show that a patient-derived antibody, AtS13, has a strong resemblance to PVIgG in its effects on adhesion dissociation. Importantly, our study also shed light on anti-non-Dsg antibodies, such as anti-TPO, which have not been known to link tightly with PV yet. Collectively, our quantitative image analysis framework can be expected to facilitate the development of entirely new class of PV diagnostic tools.

## MATERIALS AND METHOD

### Cell culture

HaCaT cells, an immortalized human keratinocyte cell line, were cultured in a low-calcium medium. This medium was prepared from calcium-free DMEM (ThermoFisher Scientific, Cat. No. 21068028) and supplemented with 10% fetal bovine serum (FBS) which contains calcium, 1 % GlutaMAX (ThermoFisher Scientific, Cat. No. 35050061), and 1% penicillin-streptomycin. The cell cultures were maintained at 37°C in a 5% CO_2_ atmosphere to ensure continuous cell division.

### Antibody treatment

Once the cells reached 80% confluency, they were switched to a calcium-supplemented medium with 1.8 mM calcium overnight before undergoing antibody treatment. All antibody treatments were conducted in the calcium-supplemented medium, using antibody concentrations of 2 μg/mL and 10 μg/mL with exposure times of 4 hours and 24 hours. PX4-4 and PX4-3, provided by Dr. Aimee Payne (Columbia University), are monoclonal antibodies derived from the single-chain variable-region fragment (ScFv) of antibodies isolated from a patient suffering from mucocutaneous PV. The anti-thyroid peroxidase (anti-TPO) antibody (NBP1-50811, Novus Biologicals) targets thyroid peroxidase, an enzyme crucial for the production of thyroid hormones. AtS13, developed by the Sinha group at the University at Buffalo, is a PV patient-derived anti-human monoclonal antibody. It shows significant homology to anti-TPO antibodies and to antibodies targeting desmosome proteins. PVIgG represents polyclonal antibodies purified from the serum of a PV patient, containing high levels of anti-Dsg3 antibodies and only trace amounts of anti-Dsg1 antibodies. The anti-Dsg3 antibody AK23 (D219-3, MBL) specifically targets Dsg3, a crucial component of cell-cell adhesion in epithelial cells, predominantly affected in PV. We also used anti-human HLA-ABC (555551, BD Biosciences) as a control antibody that is not expected to bind to cell adhesion structures.

### Immunostaining

Antibody-treated HaCaT monolayers were washed three times with phosphate-buffered saline (PBS, Thermo Fisher). The cells were then fixed for 10 minutes with 4% paraformaldehyde for Dsg3, E-cad, and F-actin staining, or for 5 minutes with ice-cold methanol-acetone (1:1) for IF staining. For RhoA staining, cells were fixed using 10% trichloroacetic acid (TCA, Sigma-Aldrich, T6399-5G) for 15 minutes at room temperature. Following fixation, the cells were permeabilized with 0.1% Triton X-100 for 5 minutes. To block nonspecific binding, the monolayers were incubated for 1 hour in a blocking buffer composed of 1% bovine serum albumin (BSA) (37520, Thermo Fisher) and 22.52 mg/mL glycine (410225, Millipore Sigma) in DPBST (Dulbecco’s phosphate-buffered saline with 0.1% Tween 20) (P7949, Millipore Sigma). This was followed by three 5-minute washes with PBS. For immunofluorescence staining, primary antibodies, including anti-Dsg3 (1:200, Abcam), anti-E-cadherin (1:200, Cell Signaling Technologies), anti-RhoA 26C4 (1:200, Santa Cruz), and Pan-Keratin C11 (1:200, Cell Signaling Technology), were applied and incubated for 1 hour. The cells were then exposed to secondary antibodies Alexa Fluor 647 (1:100, Thermo Fisher), Alexa Fluor 488 (1:100, Thermo Fisher), and Alexa Fluor 594 (1:100, Thermo Fisher) for 1 hour. F-actin was labeled using Alexa Fluor 488 Phalloidin (1:100, Thermo Fisher) with a 30-minute incubation. After staining, the monolayers were washed three additional times with PBS for 5 minutes each, then imaged following standard protocols.

### Image processing and quantification

Each microscope image was divided into four sub-images to ensure that the heterogeneity of the sample was adequately captured. To quantify texture, we focused on regions with sufficient signal, omitting empty areas such as those belonging to the cell nucleus. This was particularly important for proteins like E-cadherin, which are primarily concentrated at cell boundaries.

To identify the regions of interest (ROI) for calculating texture parameters, the sub-images were further divided into 100×100 pixel segments. These segments were screened for adequate signal before analysis. Specifically, a threshold was set at 5% of the maximum image intensity. If fewer than 80% of the pixels in a segment fell below this threshold, the segment was included in the analysis. Segments failing to meet this criterion were discarded. The texture parameters for each sub-image were calculated by averaging the parameters from the segments that were not discarded due to low signal.

Normalization was applied to the segments before calculating texture parameters to avoid the influence of image intensity on the results. This step was necessary to account for slight variations in staining process that could otherwise introduce bias unrelated to the antibody effects. Normalization involved dividing pixel intensities by the maximum intensity within each segment, effectively scaling all intensities between 0 and 1, which mitigated variations caused by staining. The texture parameters calculated for the images included Correlation, Entropy, Contrast, Homogeneity, Energy, Mean, and Standard Deviation. These parameters, derived from the Gray-Level Co-Occurrence Matrix (GLCM) analysis, provide important quantitative measures that reflect different aspects of the image’s texture.

*Correlation* measures how correlated a pixel is with its neighbors over a specified distance. This parameter gives insight into the linear dependencies of gray levels within an image, with higher values indicating stronger correlation between pixel pairs. *Entropy* is a measure of randomness or complexity in the intensity distribution of the image. A high entropy value indicates that the image has a complex, unpredictable texture with a wide range of pixel intensity values. *Contrast* refers to the local intensity variation between a pixel and its neighboring pixels. *Homogeneity* evaluates the similarity of pixel values in the local area. High homogeneity values correspond to images with more uniform intensity distributions, where pixel values are more similar to each other. *Energy* represents the uniformity of the pixel intensity distribution, calculated as the sum of squared pixel values in the GLCM. High energy values indicate that the image is dominated by certain pixel intensities, suggesting a repetitive and uniform texture. *Mean* is the average pixel intensity within a region of interest and *Standard Deviation (Std)* quantifies the spread of intensity values around the mean.

Together, these parameters provide a comprehensive view of the texture and structural features within the image, capturing both the intensity variations and the spatial relationships between pixels. In addition to these texture parameters, two additional metrics were calculated for proteins F-actin and intermediate filaments (IFs), which exhibit fibrous structures. These metrics were isotropy level (Iso) and circular variance (CV). The Iso parameter reflects the relative area of the image lacking strong fibrous structure, while CV quantifies the dispersion of fibers in 2D, with values ranging from 0 (indicating alignment in a single direction) to 1 (indicating complete dispersion). In mathematical terms, the CV of a set of vectors, 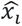, with direction cosines [*l_i_*, *m_i_*] (*i* = 1, …, *n*), is calculated as follows:

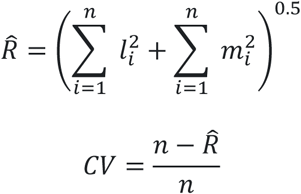

If the vectors represent fibers, another set of vectors with the opposite directions relative to the given vectors, i.e., 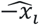, can equally represent the same fibers. Therefore, the above equation is modified to account for this property. The detailed calculation methods for these parameters are described in previous publications (34,35).

### Random Forest analysis

To classify antibodies based on quantitative image parameters, we implemented a Random Forest algorithm in R. The dataset was split into training and testing sets, with 80% of the data used for training and the remaining 20% for testing. To increase the robustness of the analysis, the antibodies were not separated based on dose or treatment time. This approach allowed the model to be trained and evaluated across a range of dosing regimens and treatment durations, reducing reliance on individual imaging sessions and enhancing the generalizability of the results. The training set was divided using stratified sampling to preserve the distribution of antibody classes. Specifically, we applied a five-fold cross-validation approach to optimize the model parameters and ensure robustness, reducing overfitting by averaging the results over multiple training subsets.

The Random Forest model was built using 200 decision trees, where each tree was trained on a bootstrapped sample of the data. At each node split, a random subset of features was selected, ensuring that the model did not overly rely on any particular variable. Following model training, we calculated the importance of each variable to assess which features contributed the most to the classification task. The importance scores were derived by measuring the decrease in classification accuracy when a specific variable’s values were permuted, effectively disrupting the relationship between the feature and the output. This enabled us to rank the variables and present their relative importance in the model.

The performance of the model was evaluated using a confusion matrix, which displayed the number of true positives, false positives, false negatives, and true negatives for each antibody class. This matrix provided a quantitative overview of the model’s classification power, highlighting the frequency of correct predictions versus errors. To further investigate the misclassifications, we identified pairs of antibodies that were frequently confused by the model. We filtered for misclassifications occurring more than twice and constructed a similarity graph based on these misclassifications. In this graph, each antibody is represented as a node, and edges between nodes represent frequent misclassifications. The weight of each edge corresponds to the frequency of misclassification between the two antibodies. The resulting graph effectively displayed how similar certain antibodies were, based on the model’s inability to distinguish them, offering insights into possible biological similarities affecting the classification.

The pathogenicity score for each antibody was defined to quantify its effectiveness relative to a control. This score integrates the impact of various parameters, weighted by their importance, and compares each antibody’s performance to that of a control antibody. Specifically, each parameter’s importance was determined based on its contribution to the Random Forest model. The importance values, reflecting the relative impact of each parameter on classification accuracy, were used to weight the parameters. For each parameter, the absolute difference between the median value of the parameter for an antibody and the median value for the control was computed. This difference was then normalized by dividing by the control median value to account for the relative change in parameter value. Finally, the weighted differences for all parameters were summed to yield the pathogenicity score for each antibody. This score reflects the extent to which the antibody deviates from the control in terms of parameter values, weighted by their importance.

In mathematical terms, the pathogenicity score for each antibody can be expressed as:

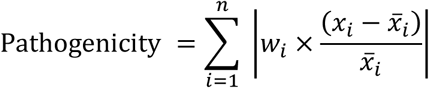

where *w_i_* is the weight of parameter (i) from the importance analysis, *x*_{*i*}_ is the median value of parameter (i) for the antibody, and {*x̄*}*_i_* is the median value of the same parameter for the control group. This approach allows for a comprehensive evaluation of each antibody’s pathogenicity by integrating multiple parameters into a single score, providing insights into the relative effectiveness of each antibody compared to the control.

### Statistical analysis

All statistical analyses were performed using RStudio (version 2023.09.1+494). A two-tailed t-test was employed to compare the mean texture values between antibody-treated and control groups, allowing for the identification of significant changes in texture features. Pearson’s correlation analysis was used to assess the relationships between texture parameters derived from immunofluorescence images, with the resulting correlation matrix offering insights into the interdependencies among these features across different antibody treatments. All statistical tests were considered significant at p<0.05.

## RESULTS

### Altered distribution of Dsg3 and E-cadherin serves as potent marker for PV pathogenicity

Using quantitative image texture features, we investigated the distribution patterns of Dsg3 and E-cadherin in keratinocytes. Cell monolayers were exposed to various PV antibodies at concentrations of either 2 µg/ml or 10 µg/ml for durations of either 4 or 24 hours. This experimental design allowed us to evaluate the effects of both dose and exposure time on the remodeling of these adhesion molecules. Immunofluorescence images revealed antibody-induced changes in the distribution pattern of Dsg3 within the cell monolayer. Certain changes, such as localized dispersion, were qualitatively evident, as indicated by the white arrows in **Fig. 1A**. A quantitative analysis allowed us to detect more subtle variations, which are detailed in **Fig. 1C-J**. The analyzed quantitative image features included entropy, correlation, standard deviation of pixel intensity, mean pixel intensity, contrast, homogeneity, and energy, collectively describing the texture of the images. Please refer to the Methods section for definitions of these parameters. Representative images of all treatment groups can be found in Supplementary Information Fig. S1.

**Figure 1.**
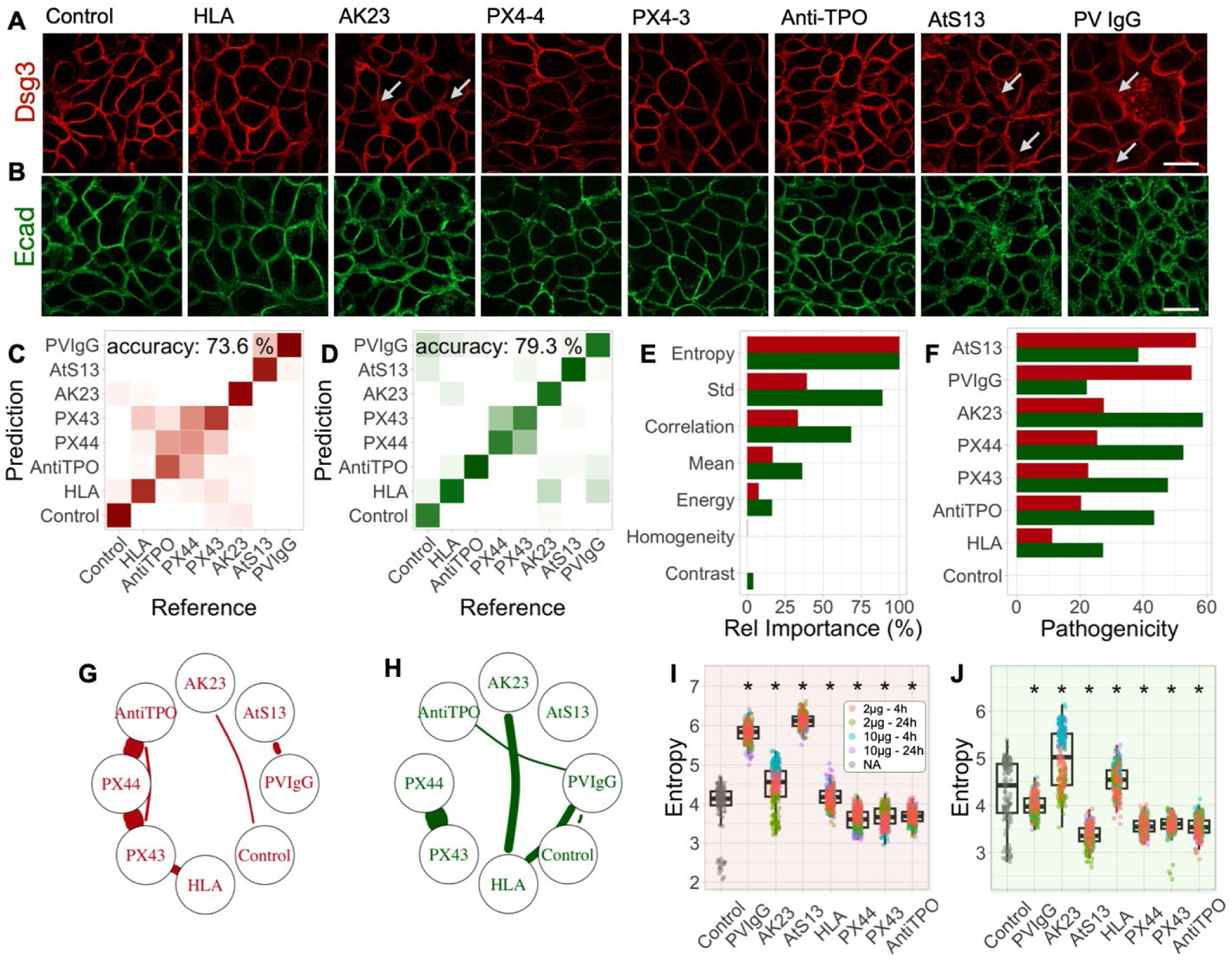
Influence of PV antibodies on immunofluorescence images of Dsg3 (red) and E-cadherin (green). A and B: Representative images for the 10 µg/ml – 24h group. The scale bar represents a distance of 25 µm. C and D: Confusion matrices resulting from the random forest analyses. A minimum of 60 images per condition were fed into the model. E: Relative importance of different image quantification parameters. F: Pathogenicity score of different antibodies. G and H: Network graphs resulting from the confusion matrices. Misclassified antibodies are connected with thicker lines. I and J: Variation of the most important parameter across different treatment groups. *Indicates a statistically significant difference (p<0.05) when the antibody group is compared to the control.

We employed a Random Forest (RF) analysis to classify the images based on quantitative texture parameters. RF algorithms are particularly advantageous due to their resistance to overfitting, accuracy with small training sets, and ability to efficiently handle large datasets. This methodology also enabled us to identify the most influential image parameters for classification. **Fig. 1C** presents the confusion matrix derived from the RF analysis. Overall, we achieved an accuracy of 73.6% in distinguishing distinct groups based on Dsg3 staining images. The model did not receive information about the dose and treatment time, and all images related to a specific antibody were treated equally. Anti-HLA antibody, which does not bind to desmosome was used as a control and had negligible effects on Dsg3. PX4-3 and PX4-4 exhibited similar effects to each other. PX4-4 is the non- to mildly-pathogenic version of the monoclonal anti-Dsg3 antibody; PX4-3 targets the extracellular domain of Dsg3 (36). These factors decreased the overall accuracy of the classification. **Fig. 1E** illustrates the relative importance of various quantitative parameters derived from the RF analysis, which indicates the contribution of each parameter to the model’s predictive performance. Among these, Entropy emerged as the most critical parameter for image classification, followed by Standard Deviation, Correlation, Mean, and Energy. Homogeneity and Contrast did not provide additional valuable insights regarding the treatment groups.

The pathogenicity score in **Fig. 1F** is a metric that quantifies the variations in parameters between the treated and control groups, considering the importance of each parameter as determined by the RF analysis. Notably, the antibodies AtS13, PVIgG, and AK23 exhibited the highest pathogenicity scores, indicating their significant influence, whereas the anti-HLA control displayed the lowest pathogenicity score in modulating the distribution of Dsg3. The similarity graph (**Fig. 1G**), based on the confusion matrix, illustrates the relative impact of antibodies on Dsg3. Antibodies that are frequently mistaken for each other are connected by thicker lines. The graph indicates that PX4-4 exhibits similar effects to both anti-TPO and PX4-3, while other similarities are comparatively weaker. The most significant quantitative imaging parameter, here Entropy, for various antibodies is shown in **Fig. 1I**. Groups that significantly differ from the control (p<0.05) are denoted with asterisks. Entropy was significantly elevated compared to the control in several groups—especially in AtS13, PVIgG, and AK23, the three antibodies with the highest potencies. An increase in Entropy might suggest that the distribution of Dsg3 in cells has become more disordered after treatment, indicating that Dsg3 proteins are less uniformly localized. Our immunofluorescence imaging analysis emphasizes that autoimmune antibodies distinctly alter the distribution patterns of Dsg3, suggesting potentially different mechanisms of action.

Next, we evaluated the changes induced by PV antibodies in E-cadherin distribution. Initial qualitative observations indicated that similar to Dsg3, E-cadherin displayed varied distribution patterns across different antibody treatments (**Fig. 1B**, Fig. S2). Employing a similar methodological framework, we performed an RF analysis on E-cadherin-stained images. This analysis yielded a notable accuracy of 79.3%, emphasizing the model’s robust capability in differentiating effects based on textural E-cadherin differences (**Fig. 1D**). Entropy, Standard Deviation, Correlation, and Mean were identified as the most important quantitative parameters for distinguishing the effects of antibodies on E-cadherin (**Fig. 1E**), which mirrors the relative importance calculated in the Dsg3 analysis. This consistency across both Dsg3 and E-cadherin reiterates the significance of these parameters in assessing the impact of antibody on adhesion molecules at the cell-cell adhesions.

AK23, PX4-4, PX4-3, and anti-TPO exhibited the highest pathogenicity, whereas PVIgG showed the lowest pathogenicity (**Fig. 1F**). Additionally, the similarity graph in **Fig. 1H** demonstrated that PX4-4 and PX4-3 exhibited the highest similarity based on quantitative texture features, often leading to confusion between the two. **Fig. 1J** illustrates the key quantitative parameter, Entropy, for the effects of PV antibodies on E-cadherin. The E-cadherin entropy, in contrast to that of Dsg3, showed fluctuation between increases and decreases for the high pathogenicity antibodies. This variability suggests that antibodies that disturb Dsg3, characterized by an increased Entropy, might have differing specificities and affinities for other targets, such as E-cadherin. These variations could lead to different degrees of disruption or stabilization. Furthermore, the downstream effects of such treatments can differ, as E-cadherin and Dsg3 are both crucial for cell adhesion but operate through distinct mechanisms. For instance, the disruption of Dsg3 by certain antibodies such as AtS13, but not by others, may activate compensatory mechanisms that lead to the upregulation or reorganization of E-cadherin-mediated adhesion. In some scenarios, this could enhance the organization of E-cadherin, resulting in decreased entropy. The findings collectively support the idea that PV antibodies induce distinguishable alterations in the distribution patterns of E-cadherin, similarly to Dsg3, proposing differential mechanisms affected by autoimmune mediators in keratinocytes.

### Cytoskeletal reorganization is less indicative of antibody pathogenicity compared to adhesion molecular re-distribution

In this section, we focus on the cytoskeletal components IF and F-actin, both of which are pivotal for cellular structure and function. Given that these proteins are fibrous in nature, we incorporated two additional parameters—Isotropy (Iso) and Circular Variance (CV)—in addition to our existing set of texture analysis metrics. For more details on these parameters, please refer to the Methods section.

Representative images of IF from cells exposed to different antibodies are shown in **Fig. 2A** and Fig. S3. Using an RF analysis across all antibody treatments, we achieved an accuracy of 70.6% in classifying changes induced by the antibodies based on images of IF (**Fig. 2C**). The parameters that emerged as most influential in this classification were Mean, Entropy, Contrast, and Standard Deviation (**Fig. 2E**). According to imaging of IF, PVIgG, AtS13, and AK23 showed the highest potencies, followed by other antibodies (**Fig. 2F**). The most similar antibodies in terms of the quantitative texture features of IF were PVIgG/AtS13 and AtS13/HLA, as depicted in the similarity graph (**Fig. 2G**). Although HLA is typically considered a secondary control in our experiments and shows low pathogenicity in other protein groups, it alters the image features of IF more than other proteins, making the images distinguishable from those treated with other antibodies; thus, based on these observations, HLA may not be an ideal control in this context. **Fig. 2I** illustrates the most important quantitative imaging parameter, here Mean, for various antibodies. Groups that differ significantly from the control (p<0.05) are marked with asterisks. Mean and Entropy—the most important parameters based on IF images—both decrease following antibody treatment in several groups including PVIgG, AK23, AtS13, and PX4-4. The reduced Entropy suggests a more predictable and orderly texture, possibly due to increased filament organization, filament aggregation, or reduced filament density or branching. The concurrent decrease in the Mean parameter in these antibodies indicates fewer high-intensity pixels, pointing to reduced filament density and potential degradation or disassembly of intermediate filaments. This likely corresponds to keratin retraction and degradation resulting from the antibody effect.

**Figure 2.**
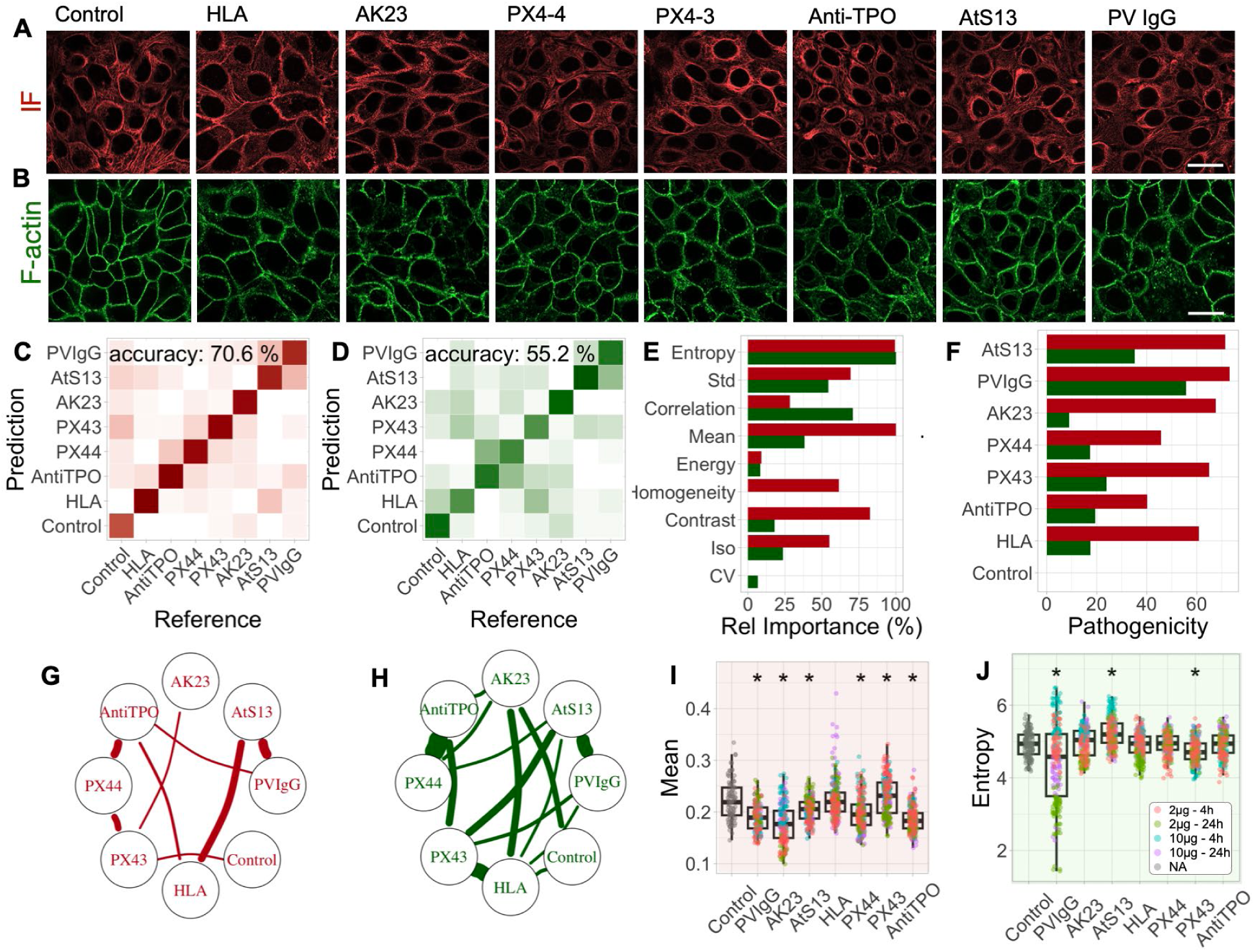
Influence of PV antibodies on immunofluorescence images of IF (red) and F-actin (green). A and B: Representative images for the 10 µg/ml – 24h group. The scale bar represents a distance of 25 µm. C and D: Confusion matrices resulting from the random forest analyses. A minimum of 60 images per condition were fed into the model. E: Relative importance of different image quantification parameters. F: Pathogenicity score of different antibodies. G and H: Network graphs resulting from the confusion matrices. Misclassified antibodies are connected with thicker lines. I and J: Variation of the most important parameter across different treatment groups. *Indicates a statistically significant difference (p<0.05) when the antibody is compared to the control.

F-actin is crucial for maintaining cell structure and mediating cellular responses to extracellular signals, making it an interesting target to study in the context of antibody-induced cellular alterations. Representative images of F-actin from cells exposed to different antibodies are shown in **Fig. 2B** and Fig. S4. The RF model achieved an overall lower accuracy of 52.2% in differentiating the treatment effects based on F-actin (**Fig. 2D**). This reduced accuracy and high confusion (**Fig. 2H**) might suggest that F-actin’s dynamic assembly and disassembly within the cell render its response to antibody treatments more complex and less predictable than the behavior of more static adhesion molecules. Regardless, the most important quantitative parameters for differentiating treatment groups were Entropy, Correlation, and Standard Deviation (**Fig. 2E**). PVIgG and AtS13 showed highest pathogenicity scores followed by PX4-3 and other antibodies (**Fig. 2F**). **Fig. 2J** illustrates the most important quantitative imaging parameter, here Entropy, for various antibodies. The impact of PV antibodies on RhoA distribution pattern is examined in Supplementary Information Fig. S5 and S6.

Collectively, the results indicate that PV-associated antibodies modulate the cytoskeletal reorganization of F-actin and IF, with IF being more distinctly influenced by various antibodies compared to F-actin.

### High dose of antibodies induces more substantial ultrastructural changes

The impact of treatment dose on the accuracy of prediction was systematically assessed by selectively removing either high-dose (10 µg/mL) or low-dose (2 µg/mL) data from the analysis (**Fig. 3A**). In the original model, the accuracy based on Dsg3 data was 73.6% (**Fig. 3D**). Upon exclusion of the high-dose data, the accuracy slightly decreased to 71.5% (**Fig. 3E**), while removing the low-dose data led to a marked improvement in accuracy, reaching 80.7% (**Fig. 3F**). This suggests that the high-dose treatment induces more significant ultrastructural changes in Dsg3, enhancing the model’s ability to predict treatment group. A similar trend was observed for E-cadherin and RhoA (SI Fig. S5, S6), where the high-dose data appeared to contribute more prominently to the model’s predictive power (**Fig. 3A**). The analysis of F-actin and IF revealed a consistent increase in prediction accuracy regardless of whether high- or low-dose data were removed, with the highest accuracy observed when low-dose data were excluded (**Fig. 3A**). This indicates that high-dose antibody treatment exerts a more substantial effect on the ultrastructure of these proteins, leading to clearer distinctions between the treatment groups. Consistent with these observations, removing the low-dose data generally increased the pathogenicity scores across most proteins, reinforcing the idea that high-dose treatments induce more pronounced changes in protein structure (**Fig. 3C**). The relative importance of imaging parameters (**Fig. 3B**) remains consistent, revealing that Entropy and Standard Deviation consistently show high importance across different treatment doses for Dsg3 images. This underscores their robustness as key features responsive to antibody treatment. In contrast, features like Contrast and Homogeneity consistently demonstrate minimal impact, indicating they may be less effective in capturing the structural changes induced by the treatments.

**Figure 3.**
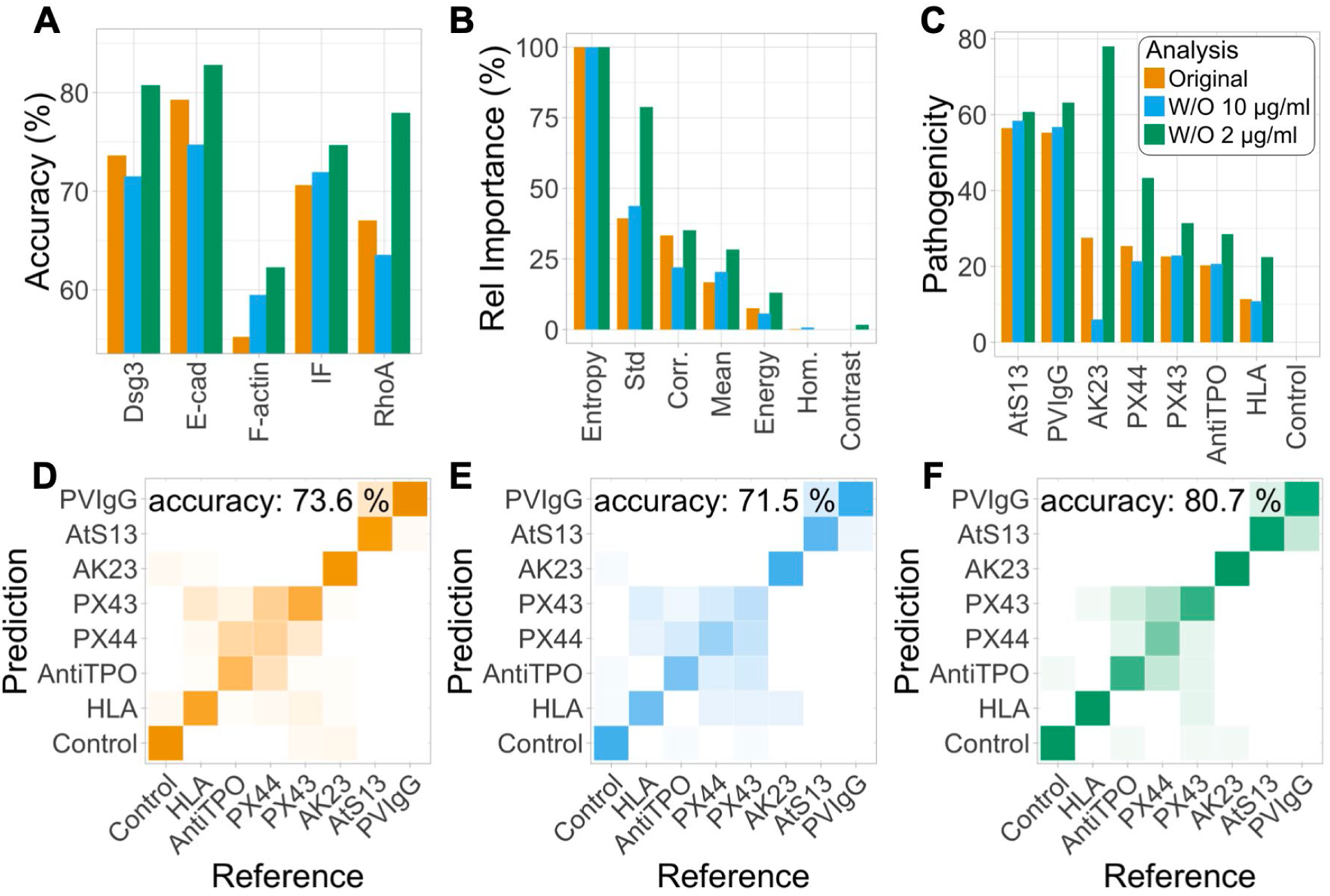
Impact of treatment dose on model predictive power. A: Prediction accuracy across various protein images. The overall accuracy, considering all antibodies, is presented. The orange color represents the complete data set from the original analysis, whereas the blue and green colors represent the data sets with the 10 µg/ml and 2 µg/ml data removed, respectively. Both the 4h and 24h treatment groups remained in the dataset and were not differentiated. B: Relative importance of imaging parameters derived from Dsg3 images. C: Pathogenicity scores calculated from Dsg3 images. D-F: Confusion matrices for different analyses based on Dsg3 images.

The mixed influence of treatment duration (Fig. S7) on prediction accuracy suggests that the temporal dynamics of ultrastructural changes vary among proteins. The overall increase in pathogenicity scores (Fig. S7) in the longer treatment group implies that prolonged exposure generally induces more pronounced changes in protein structure, consistent with literature.

### Collective remodeling of cell-cell adhesion marks antibody pathogenicity and correlation

In the previous sections, we explored how various texture measurements can be used to quantify changes in the distribution of critical adhesion molecules within cells. Building on this foundation, we now shift our focus to evaluate the interdependencies among these changes — specifically, whether alterations in certain proteins are associated with modifications in others. To systematically analyze these potential correlations, we aggregated the data based on antibody dose, exposure duration, and treatment type. We then averaged the data for all samples within each group to allow for pair-wise comparisons. Following data aggregation, we assessed the correlation between all possible pairs of parameters, computing both the correlation coefficients and the statistical significance of these correlations as depicted in **Fig. 4 A-D**.

**Figure 4.**
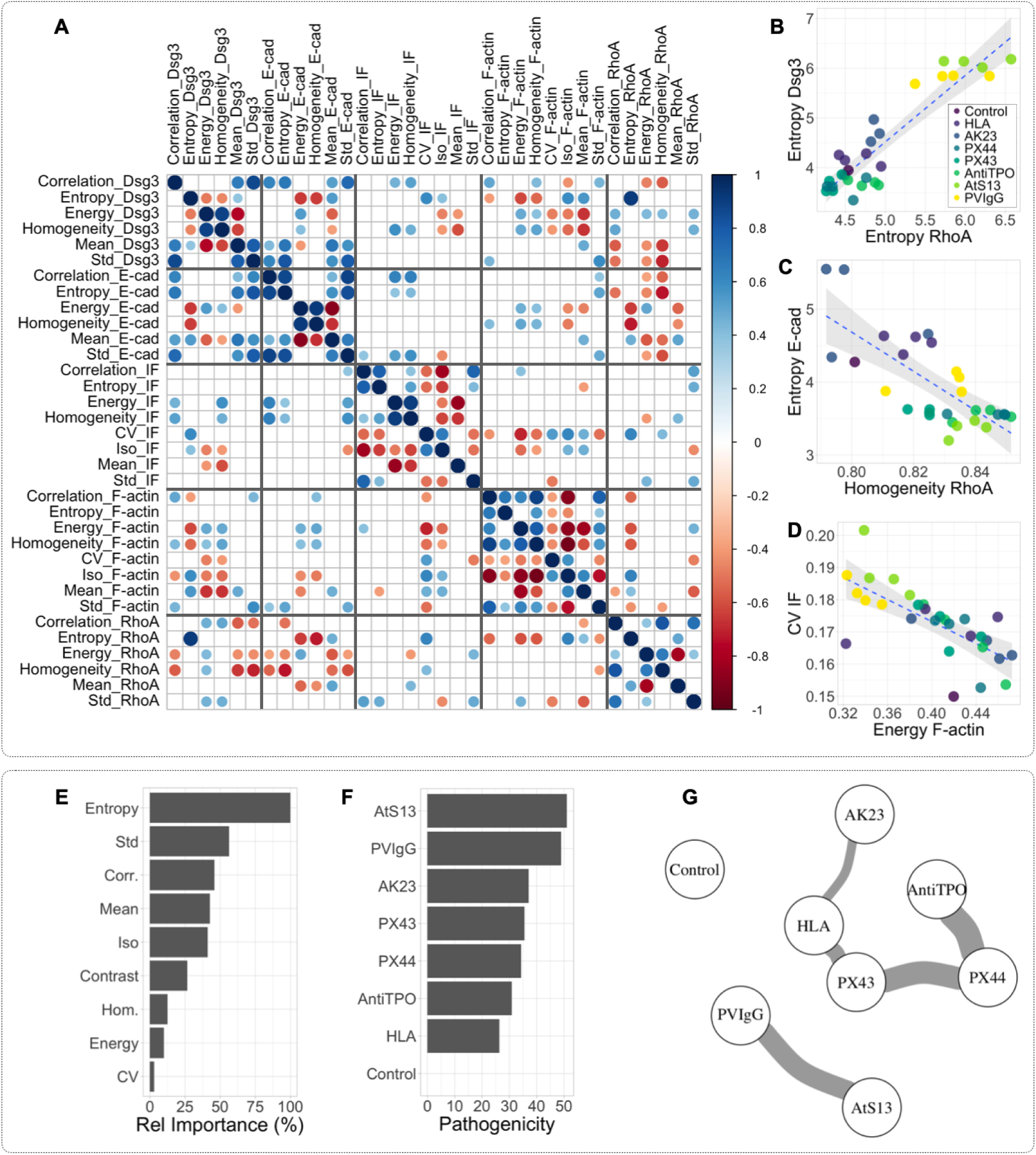
Correlation between all possible pairs of parameter-AB groups and integrative analysis. A: Correlation matrix showing all possible pairs of parameters and antibody groups, highlighting significant correlations (p<0.05). Non-significant correlations are not shown. The data are aggregated based on antibody dose, exposure duration, and treatment type. Each cell in the matrix is color-coded to represent correlation strength and direction, where blue signifies a positive correlation and red indicates a negative correlation. The size of each dot correlates with the strength of the correlation. B-D: Scatter plots illustrating representative correlations from the matrix. Colors are used to differentiate antibodies, but the dose and treatment duration are not distinguished. E: Overall relative importance of imaging parameters, averaged across all proteins and weighted by classification accuracy. F: Overall pathogenicity scores. G: Overall antibody similarity graph.

Our analysis revealed several significant correlations across different pairs. Notably, the strongest and most frequently observed correlations were between Dsg3 and E-cadherin, as well as between Dsg3 and RhoA. This pattern of correlation not only supports the hypothesis of interconnected functional pathways but also hints at possible causal relationships facilitated by antibody interactions. The pronounced correlation between Dsg3 and RhoA suggests a potential cascade effect wherein the binding of antibodies to Dsg3 may lead to downstream alterations affecting RhoA expression. This could imply that disruptions in Dsg3 integrity due to antibody binding led to cytoskeletal rearrangements mediated by RhoA, which is known for its role in regulating cellular shape and motility. Additionally, the significant correlation with E-cadherin could indicate that alterations in Dsg3 might expose or otherwise affect E-cadherin, potentially influencing cell adhesion properties and intercellular interactions. For instance, upon antibody binding to Dsg3, one might observe subsequent effects such as an increase in RhoA activity, which in turn could modulate the organization and stability of actin filaments, impacting cellular mechanics and adhesion dynamics. This might further cascade to affect the stability and exposure of E-Cadherin on the cell surface, altering cell-cell adhesion dynamics.

Given the significant correlations between remodeling processes, it is clear that a holistic approach is informative. By integrating analyses across all proteins, we can gain a more comprehensive understanding of cellular remodeling dynamics. **Fig. 4E** illustrates the overall relative importance of imaging parameters, weighted by the accuracy of the RF classification for each protein. This plot identifies the most influential parameters, with Entropy and Standard Deviation emerging as the top contributors. These findings are consistent with our analysis of individual proteins, further underscoring the significance of these parameters in distinguishing cellular states. **Fig. 4F** shows the overall pathogenicity score considering all proteins, again weighted by the RF accuracy and averaged across groups. This provides a holistic view of the treatment effects, revealing that certain antibodies, particularly the patient-derived antibodies AtS13 and polyclonal PVIgG, have a pronounced impact on cellular remodeling processes, more so than non-patient-derived monoclonal antibody sources. Finally, **Fig. 4G** illustrates a similarity graph combining all imaging data, showcasing the relationships between different treatment conditions. This network reveals clusters and connections that suggest potential common pathways or interactions affected by the antibody treatments.

Collectively, these findings underline the complex interplay between different adhesion-regulating molecules in response to specific antibody treatments. By integrating these relationships through advanced data analysis and imaging techniques, we gain a more comprehensive understanding of the pathophysiological mechanisms underpinning disorders like PV. This integrative approach underscores the importance of considering collective changes in protein distributions rather than isolated events, offering insights into potential therapeutic targets and disease dynamics.

### Texture features differentiate the responses of Dsg3 and IF to cyclic and static stretching following AK23 treatment

To showcase the predictive power and broad applicability of our data-driven approach, we examined the effects of different mechanical stretching protocols on Dsg3 and IF image features, using a dataset from reference (13). Representative images used are shown in Fig. S8. The six experimental groups comprised a control, cyclic stretch (CS), static stretch (SS), AK23 antibody treatment, AK23+CS, and AK23+SS. This approach builds on our previous work, where we established that cyclic stretch was more effective than static stretch in stabilizing cell-cell adhesion, mitigating the dissociative effects of anti-Dsg3 antibody AK23.

The analysis of Dsg3 texture features from the immunofluorescence images, presented in **Fig. 5A-C**, reveals that static stretch did not cause significant changes to Dsg3 distribution. This suggests that static stretch has minimal impact on Dsg3 organization in the control group. In contrast, cyclic stretch led to significant alterations in 2 of the 3 image parameters, i.e. entropy and correlation, indicating that Dsg3 is more responsive to cyclic mechanical forces. The AK23 antibody treatment resulted in significant changes in all texture parameters, supporting the use of these features to robustly characterize antibody-induced dissociation. Notably, the combination of AK23 and cyclic stretch further deviated the texture parameters from the control, while AK23 with static stretch brought the parameters closer to control levels. Under these experimental conditions static stretching may offer a more protective effect on Dsg3 integrity, in contrast to our previous results (13). It is important to acknowledge that the quality of Dsg3 images in the previous study, which lacked the use of a confocal microscope, may have contributed to differences in the conclusions, as lower-quality images could obscure subtle effects.

**Figure 5.**
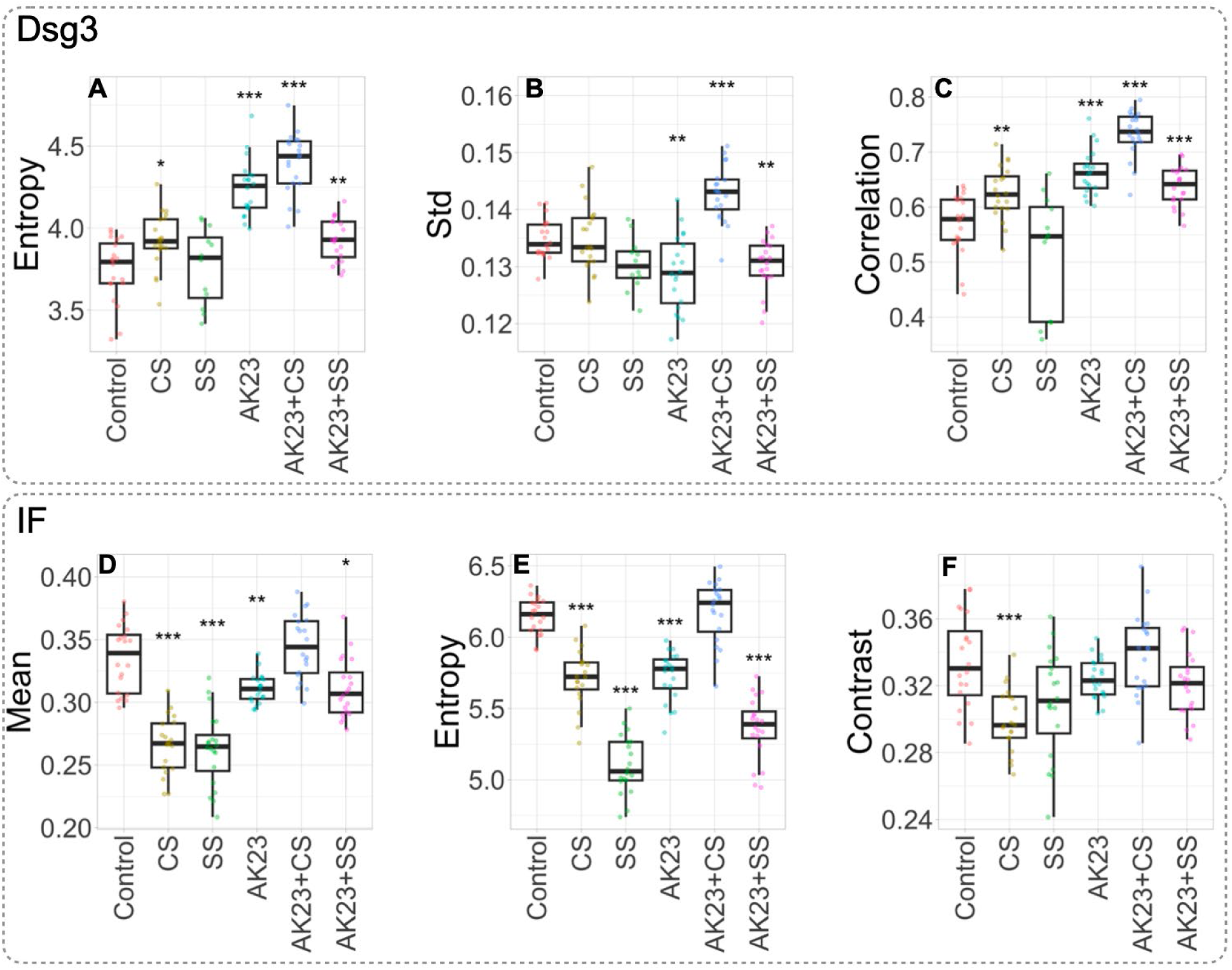
Quantification of texture features in Dsg3 and IF images under various mechanical and antibody treatment conditions. Boxplots representing the distribution of texture parameters for Dsg3 (A-C) and IF (D-F) under six experimental groups: Control, Cyclic Stretch (CS), Static Stretch (SS), AK23 antibody treatment, AK23 + CS, and AK23 + SS. The texture features quantified include Entropy (A, E), Standard Deviation (Std) (B), Correlation (C), Mean (D), and Contrast (F). Statistical significance is indicated as follows: *p<0.05, **p<0.01, **p<0.001.

For the IF texture features shown in **Fig. 5D-F**, both cyclic and static stretch significantly affected IF structure, with one exception. Among the texture metrics, Entropy was particularly sensitive to the type of mechanical loading, allowing differentiation between the effects of cyclic and static stretch on IF organization. The AK23 treatment caused significant changes in 2 out of 3 texture parameters, confirming its disruptive influence on IF organization. Interestingly, in parameters affected by AK23 treatment—Mean and Entropy—cyclic stretch, combined with AK23, restored the texture features to control-like values, effectively reversing the damage induced by AK23. In contrast, the combination of static stretch and AK23 did not fully restore these features, resulting in values that remained significantly different from the control. This observation aligns with our previous conclusions that cyclic stretch has a protective and restorative effect against AK23-induced dissociation, while static stretch does not. The higher image quality of the IF data could partly explain the more robust findings in the IF analysis.

## DISCUSSION

Our study uses a data-driven approach to analyze images of keratinocyte monolayers obtained from in vitro experiments, and we demonstrate that the disruption of cell-cell adhesion molecules and the cytoskeleton remodeling can serve as robust markers for antibody pathogenicity for PV. Specifically, we showed that the patterns of Dsg3 distribution at cell-cell contacts accurately distinguish Ab-treated groups from controls. While staining of Dsg3 in control keratinocytes show tightly packed Dsg3 at the cell-cell borders, keratinocytes after antibody treatment have a dispersed Dsg3 distribution and non-continuous cell-cell borders. These features are captured in the image analysis, most dominantly by a change in entropy from control samples. A high entropy value indicates that an image has a complex and unpredictable texture. In the context of biological images, entropy may be indicative of complex organization or heterogeneous protein expression. Similar accuracy was achieved using the patterns of E-cadherin distribution, which was also significantly distorted with PV antibody treatment.

Cytoskeleton remodeling after PV antibody treatment, particularly the remodeling of keratin, can also be captured with image analysis with high accuracy. This is not surprising given that the immediate impact of desmosome disassembly is the initiation of keratin retraction, which results in the loss of keratin in cell periphery and the change of keratin curvature at the cell periphery. These biophysical features are primarily captured using entropy parameters, along with several other parameters. Further, our analysis also matches the time course of antibody action. For instance, it has been shown that keratin retraction and Dsg3 depletion occur after 2 hours of antibody treatment (18). Our data at 4 hours also starts to show significant changes after treatment with PV-associated antibodies compared with controls. It is worth noting that the cytoskeleton remodeling data is corroborated by our analysis of RhoA imaging.

One of the main contributions of this work is that our data-driven model can reveal the pathogenicity of antibodies. Based on the overall analysis, we found patient-derived AtS13 and PVIgG to be the most pathogenic while anti-HLA antibodies, which are not expected to bind to cell-cell adhesion structures, have the lowest pathogenicity score. This pathogenic score data matches with our dissociation assay results, where we show PVIgG induces the most fragments (7). This data is also consistent with our previous findings showing that PVIgG leads to the most significant tension loss in Dsg3 molecules from FRET sensor studies compared with AK23, a monoclonal anti-Dsg3 antibody (7).

Importantly, our data also highlight the importance of anti-non Dsg antibodies in PV. Specifically, we found that anti-TPO, an antibody that is less known to be related to PV but has been found in the serum of a high percentage of PV patients (37), exhibits relatively high pathogenicity in altering proteins, such as E-cadherin. Its mechanism of action and role in PV pathology is yet to be fully explored. In addition to pathogenicity scores, analyzing image features after antibody treatment also reveals the correlation between different PV antibodies. For instance, we were able to show the similarity of PX4-3 and PX4-4 based on the analysis of Dsg3 patterns and IF remodeling. Another interesting outcome is the correlation of AtS13 antibody activity with that of PVIgG. The target of AtS13 has not been identified but is distinct from Dsg3/1.

Admittedly, the predicting power of our method is strongly dependent on the quality of immunofluorescence images. For instance, we have shown that Entropy was proved to be an effective feature on Dsg3 images from a confocal microscope for revealing adhesion dissociation. However, when applied to lower-quality Dsg3 images collected with a regular fluorescence microscope, the restoring effect of cyclic stretch cannot be predicted, suggesting a re-training of feature parameters on these sets of images. However, this restoring effect of cyclic stretch can be predicted with high-quality IF images collected using confocal microscope.

Our data driven approach to assess pathogenicity of specific antibodies within PV subgroups may be of potential use in understanding varying clinical presentations. For instance, pathogenicity scores could stratify antibody potency and pathogenicity within and across individuals who are in various clinical categorizations, such as active vs. early/late remission stages of disease, or to lesional localizations, such as mucosal vs. mucocutaneous blister distributions. This information would be helpful to clarify our thus-far opaque understanding of disease pathomechanisms, the evolution of disease, and clinical heterogeneity. This experimental approach will also be valuable to compare and contrast the effects of anti-Dsg vs. anti-non Dsg antibodies, whose functional role in lesional development has yet to be fully characterized. Pathogenicity scores of all specific PV related autoantibodies, in isolation and in combination within individual patients may pave the path towards immune specific and personalized and management strategies. In their future applications these analyses may also help to determine the in-situ effects of new and emerging therapeutics.

## ACKNOWLEDGMENTS

We acknowledge fundings from the NSF (award #1826135, #2143997), and the NIH National Institutes of General Medical Sciences (R35GM150623).

## AUTHOR CONTRIBUTIONS

All authors contributed to the writing and editing of the manuscript and have approved the final version. RY and AAS conceived the study, acquired funding, provided resources, and supervised the project. AOM conceptualized and implemented the data analysis approach, visualized the results, and wrote the first draft of the manuscript. XJ led the data collection, with support from JR, AOM, and HZ. BTS contributed to developing the data analysis ideas, and KSS and ML provided patient-derived antibodies for the study.

