## Supplemental Information for "Data-Driven Image Analysis to Determine Antibody-Induced Dissociation of Cell-Cell Adhesion and Antibody Pathogenicity in Pemphigus Vulgaris"

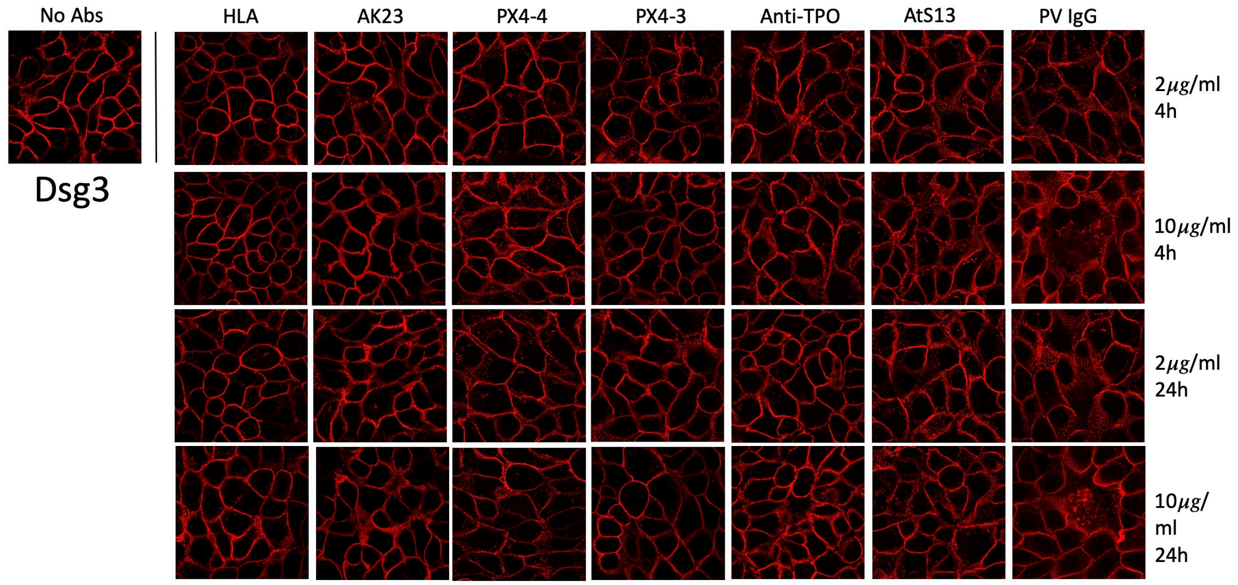


**Figure S1:** Representative images of Dsg3 in control HaCaT cells and HaCaT cells exposed to varying durations and concentrations of antibodies. Each antibody, dose, and time point were imaged in a minimum of three independent experiments. For each condition, at least 15 images were acquired, resulting in a total of at least 60 cropped images—four times the number of original images—being input into the RF model.

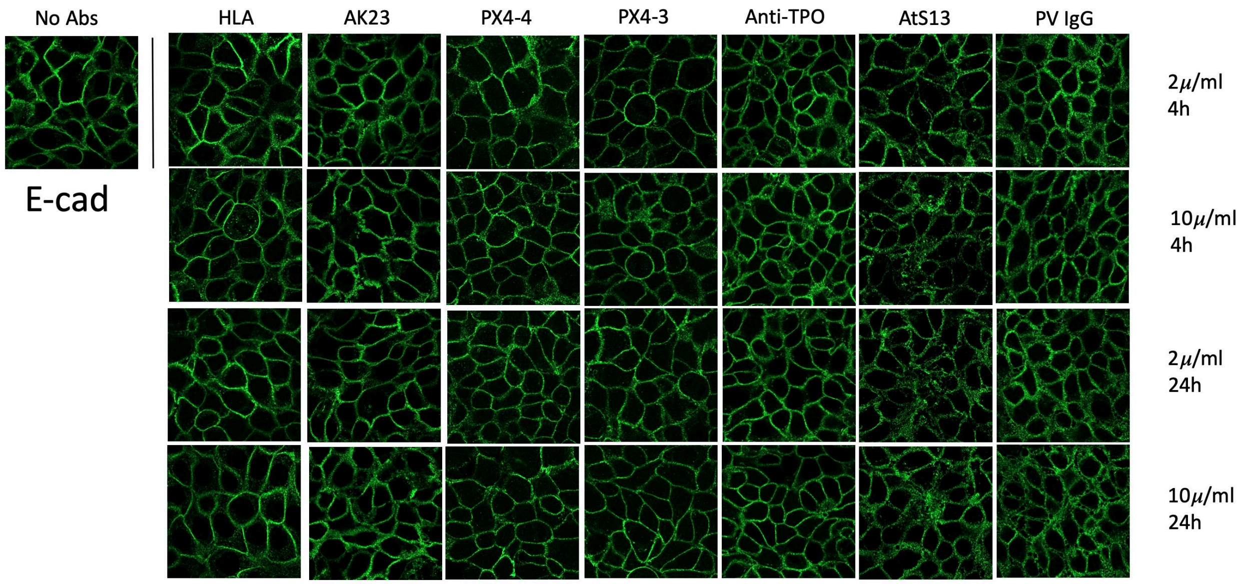


**Figure S2:** Representative images of E-cad in control HaCaT cells and HaCaT cells exposed to varying durations and concentrations of antibodies. Each antibody, dose, and time point were imaged in a minimum of three independent experiments. For each condition, at least 15 images were acquired, resulting in a total of at least 60 cropped images—four times the number of original images—being input into the RF model.

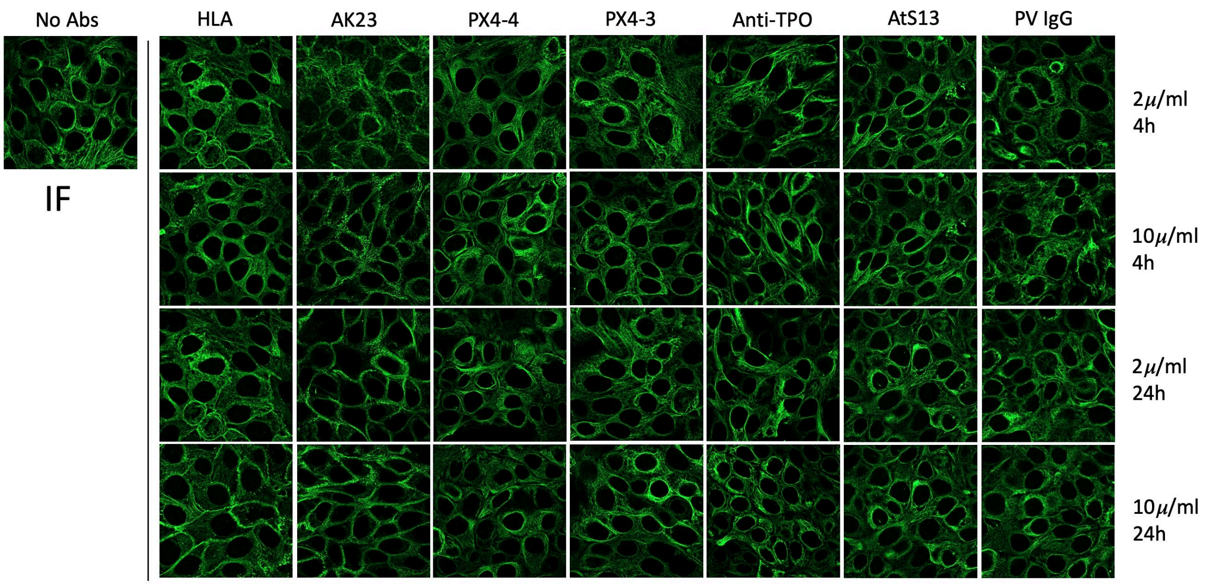


**Figure S3:** Representative images of IF in control HaCaT cells and HaCaT cells exposed to varying durations and concentrations of antibodies. Each antibody, dose, and time point were imaged in a minimum of three independent experiments. For each condition, at least 15 images were acquired, resulting in a total of at least 60 cropped images—four times the number of original images—being input into the RF model.

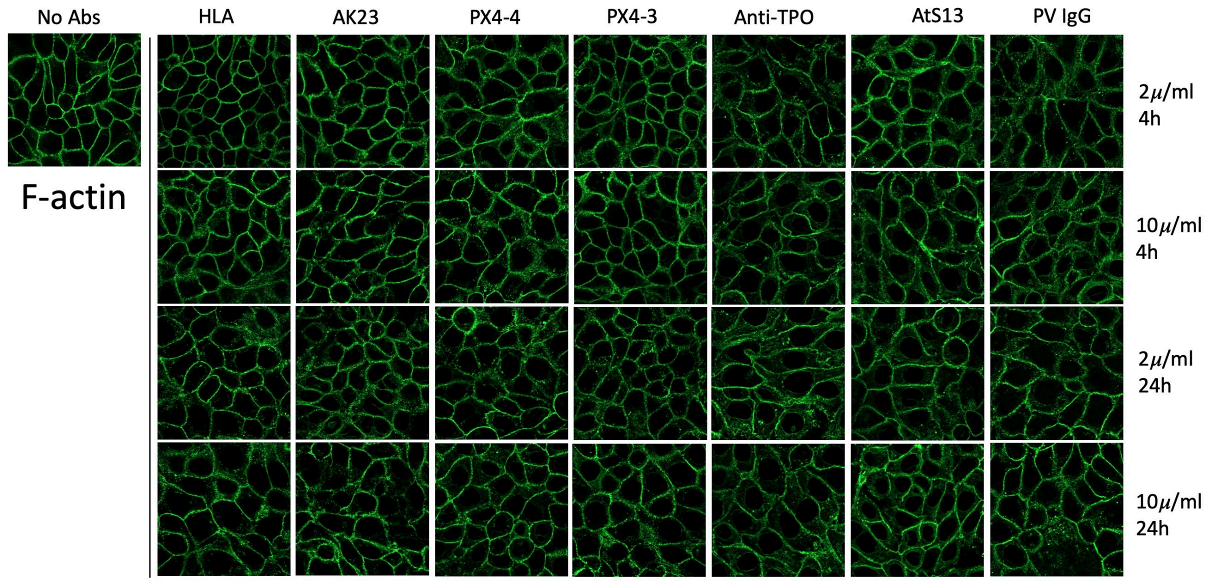


**Figure S4:** Representative images of F-actin in control HaCaT cells and HaCaT cells exposed to varying durations and concentrations of antibodies. Each antibody, dose, and time point were imaged in a minimum of three independent experiments. For each condition, at least 15 images were acquired, resulting in a total of at least 60 cropped images—four times the number of original images—being input into the RF model.

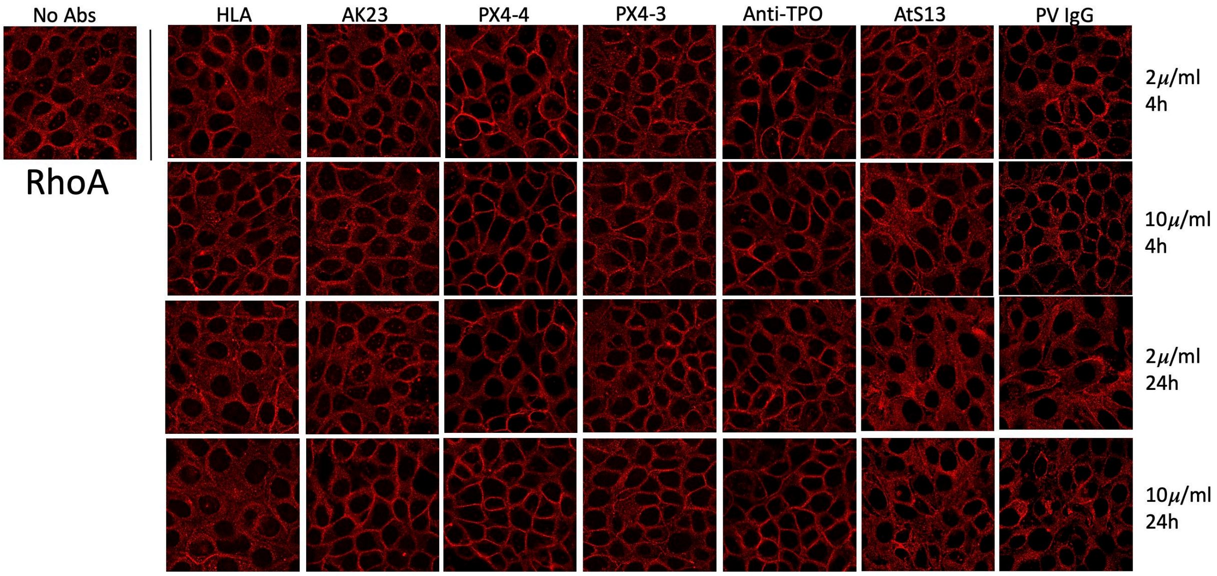


**Figure S5:** Representative images of RhoA in control HaCaT cells and HaCaT cells exposed to varying durations and concentrations of antibodies. Each antibody, dose, and time point were imaged in a minimum of three independent experiments. For each condition, at least 15 images were acquired, resulting in a total of at least 60 cropped images—four times the number of original images—being input into the RF model.

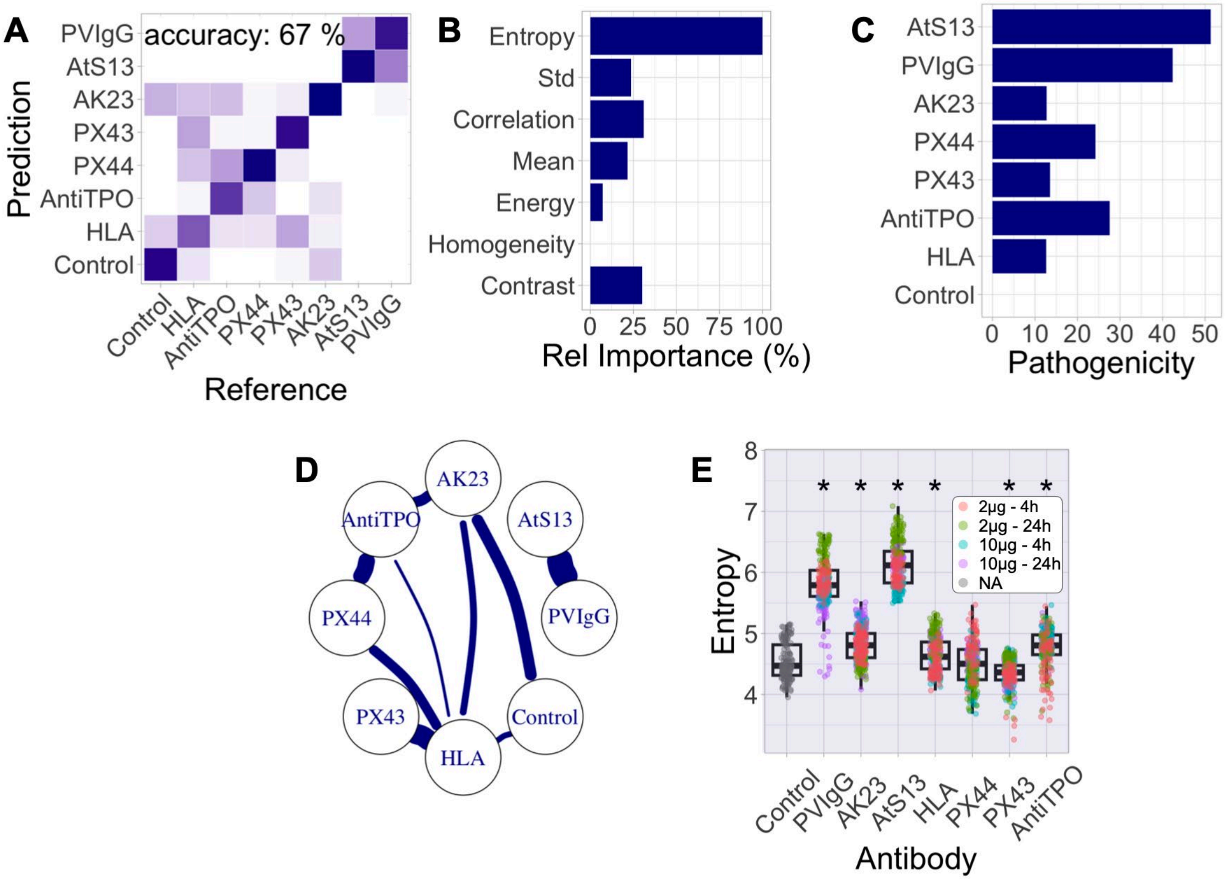


**Figure S6. Influence of PV antibodies on immunofluorescence images of RhoA.** A: Confusion matrix resulting from random forest analysis. B: Relative importance of different image quantification parameters. Entropy is the most critical parameter according to RF analysis, highlighting its importance in capturing antibody-induced alterations. C: Pathogenicity score of different antibodies. Antibodies AtS13 and PVIgG have the most significant impact on RhoA distribution, with a notable overlap in their effects. D: Network graph resulting from the confusion matrix. Misclassified antibodies are connected with thicker lines. E: Variation of the most important parameters across different treatment groups. The findings confirm that PV-associated antibodies significantly alter RhoA distribution, emphasizing the importance of these changes in PV pathogenesis and the need for targeted therapeutic strategies.

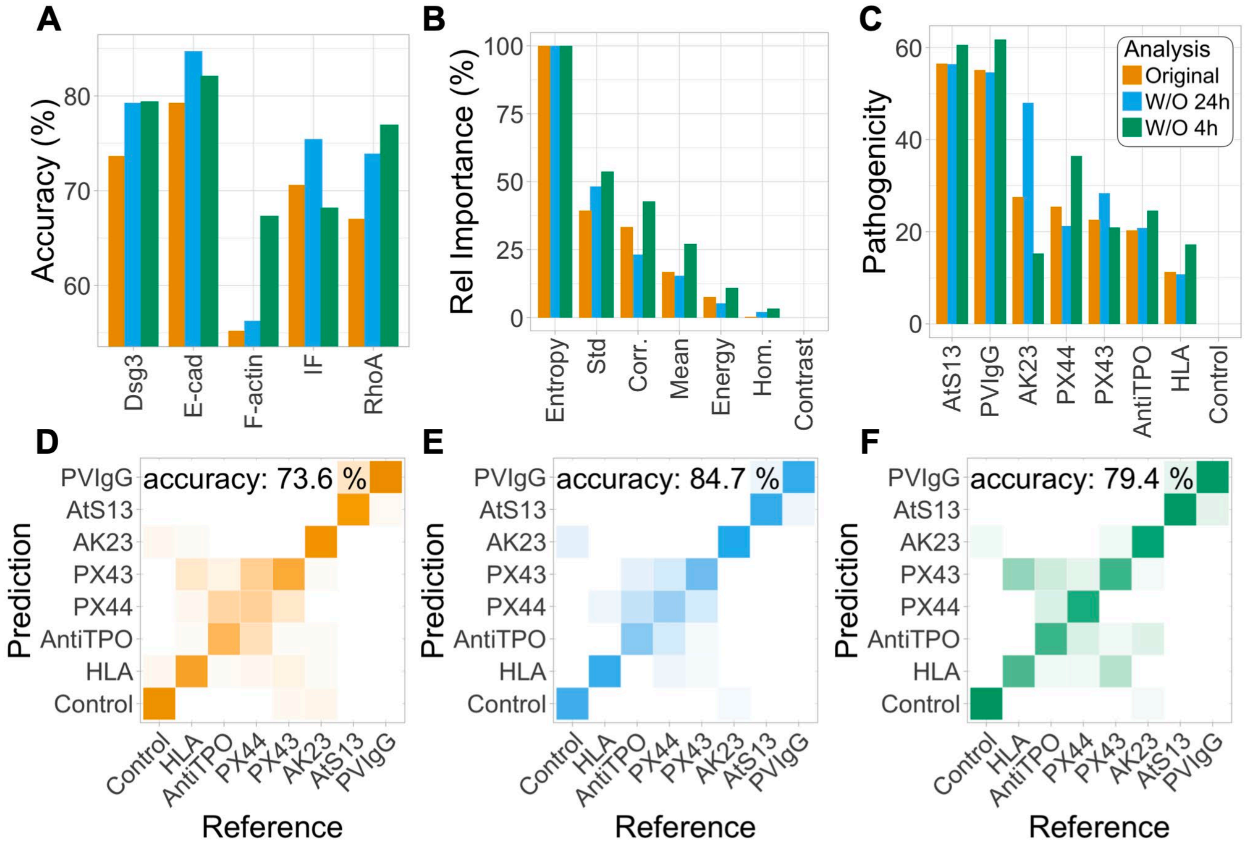


**Figure S7: Impact of treatment time on model predictive power.** A: Prediction accuracy across various protein images. B: Relative importance of imaging parameters derived from Dsg3 images. C: Pathogenicity scores calculated from Dsg3 images. D-F: Confusion matrices for different analyses based on Dsg3 images.

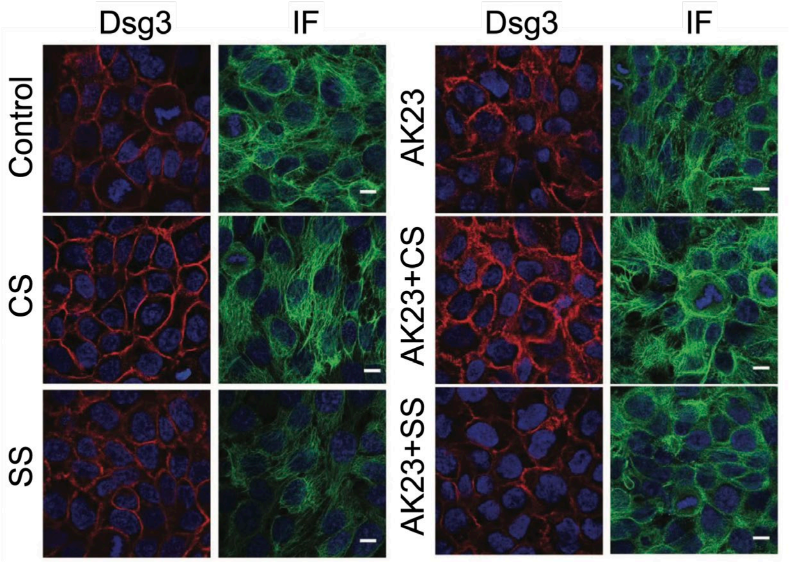


**Figure S8: Impact of cyclic stretching (CS) and static stretching (SS) on Dsg3 (red) and IF (green) following AK23 treatment (Reference 37).** Confluent keratinocytes were treated with AK23 monoclonal antibody under varying conditions of cyclic and static stretch. Subsequently, the cells were stained and imaged to examine the changes in expression and distribution of desmosome-associated proteins, including Dsg3 and IF, in response to the different treatments. Scale bar: 10 µm.
